# Neural Signatures of Prediction Errors in a Decision-Making Task Are Modulated by Action Execution Failures

**DOI:** 10.1101/474361

**Authors:** Samuel D. McDougle, Peter A. Butcher, Darius Parvin, Fasial Mushtaq, Yael Niv, Richard B. Ivry, Jordan A. Taylor

## Abstract

Decisions must be implemented through actions, and actions are prone to error. As such, when an expected outcome is not obtained, an individual should not only be sensitive to whether the choice itself was suboptimal, but also whether the action required to indicate that choice was executed successfully. The intelligent assignment of credit to action execution versus action selection has clear ecological utility for the learner. To explore this scenario, we used a modified version of a classic reinforcement learning task in which feedback indicated if negative prediction errors were, or were not, associated with execution errors. Using fMRI, we asked if prediction error computations in the human striatum, a key substrate in reinforcement learning and decision making, are modulated when a failure in action execution results in the negative outcome. Participants were more tolerant of non-rewarded outcomes when these resulted from execution errors versus when execution was successful but the reward was withheld. Consistent with this behavior, a model-driven analysis of neural activity revealed an attenuation of the signal associated with negative reward prediction error in the striatum following execution failures. These results converge with other lines of evidence suggesting that prediction errors in the mesostriatal dopamine system integrate high-level information during the evaluation of instantaneous reward outcomes.

## Introduction

When a desired outcome is not obtained during instrumental learning, the agent should be compelled to learn why. For instance, if an opposing player hits a home run, a baseball pitcher needs to properly assign credit for the negative outcome: The error could have been in the decision about the chosen action (e.g., throwing a curveball rather than a fastball) or the execution of that decision (e.g., letting the curveball break over the plate rather than away from the hitter, as planned). Here we ask if teaching signals in the striatum, a crucial region for reinforcement learning, are sensitive to this dissociation.

The striatum is hypothesized to receive reward prediction error (RPE) signals -- the difference between received and expected rewards -- from midbrain dopamine neurons (Barto, 1995; Montague et al., 1996; Schultz et al., 1997). The most common description of an RPE is as a “model-free” error, computed relative to the scalar value of a particular action, which itself reflects a common-currency based on a running average of previous rewards contingent on that action (Langdon et al., 2017). However, recent work suggests that RPE signals in the striatum can also reflect “model-based” information (Daw et al., 2011), where the prediction error is based on an internal simulation of future states. Moreover, human striatal RPEs have been shown to be affected by a slew of cognitive factors, including attention (Leong et al., 2017), episodic memory (Bornstein et al., 2017; Wimmer et al., 2014), working memory (Collins et al., 2017), and hierarchical task structure (Ribas-Fernandes et al., 2011). These results indicate that the information carried in striatal RPEs may be more complex than a straightforward model-free computation, and can be influenced by various top-down processes. The influence of these additional top-down processes may serve the striatal-based learning system by identifying variables or features relevant to the task.

To date, studies examining the neural correlates of decision making have used tasks in which participants indicate their choices with button presses or lever movements, conditions that generally exclude execution errors; as such, the outcome can be assigned to the decision itself (e.g., choosing stimulus A over stimulus B), rather than its implementation (e.g., failing to properly acquire stimulus A). To introduce this latter negative outcome, we previously conducted behavioral studies in which we modified a classic 2-arm bandit task, requiring participants to indicate their choices by physically reaching to the chosen stimulus under conditions where the arm movement was obscured from direct vision (McDougle et al., 2016; Parvin et al., 2018). By manipulating the visual feedback available to the participant, we created a series of reward outcomes that matched those provided in a standard button-pressing control condition, but with two types of failed outcomes: “Execution failures” in the reaching task, and “selection errors” in the button press task. The results revealed a strong difference in behavior between the two conditions, manifest as a willingness to choose a stimulus that had a high reward payoff, but low execution success (i.e., participants showed diminished aversion to unrewarded “execution error” trials). By using reinforcement-learning models, we could account for this result as an attenuation in value updating following execution errors relative to selection errors; in other words, when reward was withheld due to a salient execution error, participants were unlikely to decrease the value of the stimulus that they had chosen.

While this behavioral result is intuitive, the underlying neural processes are not clear. Will prediction errors in the striatum already be sensitive to the source of the error, or is the modulation of learning done through a separate top-down signal? To test this, we used fMRI to measure reward prediction errors in the striatum after both selection and execution errors. Based on our model, we hypothesized that negative prediction errors in the striatum may be weakened in the presence of salient execution failures, leading to diminished value updating.

## Methods

### Participants

A total of 24 participants were tested. The participants were fluent English speakers with normal or corrected-to-normal vision. They were all right-handed as confirmed by the Edinburgh Handedness Inventory (Oldfield, 1971). We excluded the data from four participants in the final analysis because of excessive head motion (*a priori* maximum movement threshold = 3 mm), leaving a final sample of 20 participants (11 female; age range: 18–42 years). Participants were paid $20 per hour for ~2 h of participation, plus a monetary bonus based on task performance. The protocol was approved by the institutional review board at Princeton University and was performed in accordance with the declaration of Helsinki.

### Task and Apparatus

The experimental task was a modified version of a “multi-armed bandit” task commonly used in studies of reinforcement learning (Daw et al., 2006). On each trial, three stimuli were presented, and the participant was required to choose one (Figure 1A). The participant was instructed that each stimulus had some probability of yielding a reward and that they should try and earn as much money as possible. Critically, the participant was told that each trial was an independent lottery (i.e., that the outcome on trial t-1 did not influence the outcome on trial t), and that they had a fixed number of trials in the task over which to maximize their earnings.

**Figure 1:**
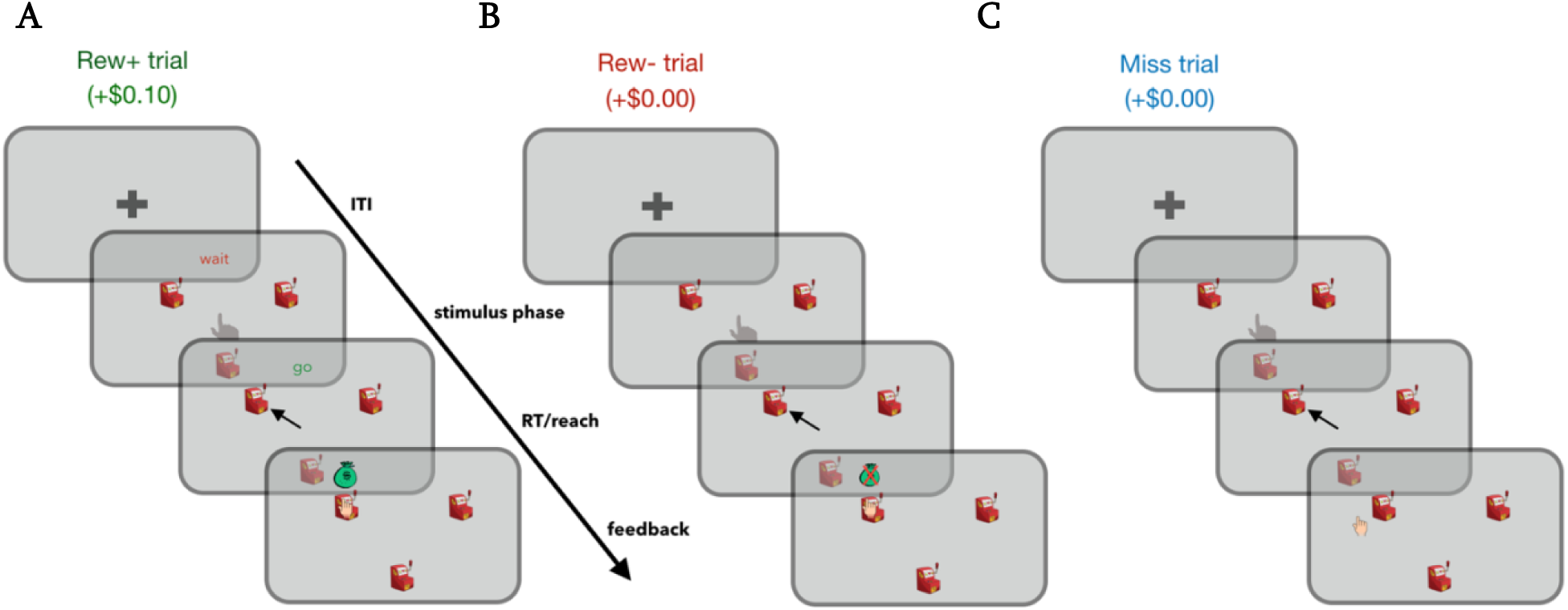
Task Design. Participants selected one of three slot machines on each trial by reaching to one of them using a digital tablet in the fMRI scanner. Three trial outcomes were possible: On Rew+ trials (A), the cursor hit the target and a reward was received; on Rew-trials (B), the cursor also hit the target but no reward was received; on Miss trials (C), the cursor was shown landing outside the target and no reward was received

In a departure from the button-press responses used in standard versions of bandit tasks, participants in the current study were required to indicate their decisions by making a wrist movement with the right hand toward the desired stimulus. The movement was performed by moving a wooden dowel (held like a pen) across an MRI-compatible drawing tablet. The tablet rested on the participant’s lap, supported by pillow wedges. The visual display was projected on a mirror attached to the MRI head coil, and the participant’s hand and the tablet were not visible to the participant. All stimuli were displayed on a black background.

To initiate each trial, the participant moved their hand into a start area, which corresponded to the center of the tablet and the visual display. The start area was displayed as a hollow white circle (radius 0.75 cm) and a message, “Go to Start”, was displayed until the hand reached the start position. To assist the participant in finding the start position, a white feedback cursor (radius 0.25 cm) corresponding to the hand position was visible when the pen was within 4 cm of the start circle. As soon as the cursor entered the start circle, the start circle filled in with white and the cursor disappeared, and the three choice stimuli were displayed along with the text “Wait” displayed in red font. The three choice stimuli were cartoons of slot machines (0.6 cm by 0.6 cm). They were presented at the same locations for all trials, with the three stimuli displayed along an invisible ring (radius 4.0 cm) at 30°, 150°, and 270° degrees relative to the origin. If the hand exited the start circle during the “Wait” phase, the stimuli disappeared and the “Go to Start” phase was reinitialized.

After an exponentially determined jitter (mean 1 s, truncated range = 1.5 s − 6 s), the “Wait” text was replaced with the message “GO!” in green font. Reaction time (RT) was computed as the interval between the appearance of the go signal and the moment when the participant’s hand left the area corresponding to the start circle. The participant had 2 s to begin the reach; if the RT was greater than 2 s, the trial was aborted and the message “Too Slow” appeared. Once initiated, a reach was considered complete when the radial amplitude of the movement reached 4 cm, the distance to the invisible ring. This moment defined the movement time (MT) interval. If the MT exceeded 1 s, the trial was aborted and the message “Reach Faster” was displayed.

The feedback cursor was turned off during the entirety of the reach. On trials in which the reach terminated within the required spatial boundaries (see below) and met the temporal criteria, reach feedback was provided by a small, hand-shaped cursor (dimensions: 0.35 cm X

0.35 cm) that reappeared at the end of the reach, displayed along the invisible ring. The actual position of this feedback cursor was occasionally controlled by the experimenter (see below), although the participant was led to believe that it corresponded to their veridical hand position at 4 cm. To help maintain this belief, the trial was aborted if the reach was > ± 25° degrees away from any one of the three stimuli, and the message “Please Reach Closer” was displayed. The cursor feedback remained on the screen for 1.5 s, and the participant was instructed to maintain the final hand position during this period. In addition to the starting circle, slot machines, and, when appropriate, feedback cursor, the display screen also contained a scoreboard (dimensions: 3.3 cm X 1.2 cm), presented at the top of the screen. The scoreboard showed a running tally of participant’s earnings in dollars. At the end of the feedback period, the entire display was cleared and replaced by a fixation cross presented at the center for an exponentially jittered inter-trial interval (mean 3 s, truncated range = 2 - 8 s).

Assuming the trial was successfully completed (reach initiated and completed in a timely manner and terminated within 25° of a slot machine), there were three possible trial outcomes (Figure 1). Two of these outcomes corresponded to trials in which the hand-shaped feedback cursor appeared fully enclosed within the chosen stimulus, indicating to the participant that they had been successful in querying the selected slot machine. On Rew+ trials (Figure 1A), the feedback cursor was accompanied by the appearance of a small money-bag cartoon above the chosen stimulus and $0.10 would be added to the participant’s total. On Rew-trials (Figure 1B), the feedback cursor was accompanied by the same money-bag overlaid with a red “X” and no money was added to the participant’s total. The third outcome consisted of “Miss” trials, in which the feedback cursor appeared outside the chosen stimulus, indicating an execution error. No money bag was presented on these trials and the monetary total remained unchanged, as in Rew-trials. Participants were informed at the start of the experiment that, like Rew-trials, no reward would be earned on trials in which their reach failed to hit the chosen target. Importantly, the outcomes for each stimulus were predetermined according to an experimenter-defined schedule (see below), and were not directly related to the actual reach accuracy of the participant.

In summary, of the three possible outcomes, one yielded a positive reward and two yielded no reward. For the latter two outcomes, the feedback distinguished between trials in which the execution of the decision was signaled as accurate but the slot machine failed to provide a payout (Rew-), and trials in which execution was signaled as inaccurate (Miss).

Unbeknownst to the participants, outcome probabilities were fixed for each target: For all three targets, the probability of obtaining a reward (Rew+) was 0.4. Targets differed in their ratio of Rew-and Miss probabilities, with each of the three targets randomly assigned to one of the following ratios for these two outcomes: 0.5/0.1 (low miss), 0.3/0.3 (medium miss), and 0.1/0.5 (high miss). In this manner, the targets varied in terms of how likely they were to result in execution errors (and, inversely, selection errors), but not in the probability of obtaining a reward. The positions of the stimuli assigned to the three Rew-/Miss probability ratios were counterbalanced across participants. Because of the fixed outcome probabilities, there is no optimal choice behavior in this task; that is, participants would earn the same total bonus (in the limit) regardless of their choices, consistent with our previous study (McDougle et al., 2016). Their behavioral strategy therefore reflected directly their attitude to the different kinds of errors.

To maintain fixed probabilities for each target, we varied whether the cursor feedback was veridical on a trial-by-trial basis. Once a target was selected (i.e., the participant initiated a reach towards the target), the outcome (i.e. Rew+, Rew-, or Miss) was determined based on the fixed probabilities. If the true movement outcome matched the probabilistically determined outcome — either because the participant hit the target on a Rew+ or Rew-trial, or missed the target on a Miss trial — the cursor position was veridical. However, if the true movement outcome did not match the probabilistically determined outcome, the cursor feedback was perturbed: If the movement had missed the target (>±3° from the center of the target) on Rew+ and Rew-trials, the cursor was depicted to land within the target. If the movement had hit the target on a Miss trial, then the cursor was depicted to land outside the target. The size of the displacement on Miss trials was drawn from a skewed normal distribution (mean 19 ± 2.3°), which was truncated to not be less than 3° (the target hit threshold) or greater than 25° (the criterion required for a valid reach), thus yielding both a range of salient errors, but also keeping errors within the predetermined bounds (values were determined through pilot testing). The direction of the displacement from the target was randomized. Given the difficulty of the reaching task (i.e., no feedback during movement, a transformed mapping from tablet to screen, small visual targets, etc.) and the strict temporal (< 1 s) and spatial (within 25° of the target) movement constraints, we expected that participants would be unaware of the feedback manipulation (see Results).

The experimental task was programmed in MATLAB (MathWorks), using the Psychophysics Toolbox (Daw et al. 2006; Brainard, 1997). Participants were familiarized with the task during the structural scan and performed 30 practice trials for which they were not financially rewarded. Participants received a post-experiment questionnaire at the end of the task to query their awareness of perturbed feedback.

## Behavioral analysis

Trials were excluded from the analysis if the reach was initiated too slowly (RT > 2 s; 0.4 ± 0.7% of trials), completed too slowly (MT > 1 s; 2.4 ± 4.5% of trials), or terminated out of bounds (Reach terminated > 25° from a target; 1.2 ± 2.0% of trials). For the remaining data, we first evaluated the participants’ choice biases: For each target, the choice bias was computed by dividing the number of times the participant chose that target by the total number of choice trials. Second, we looked at switching biases. These were computed as the probability that the participant switched to a different target on trial t given the outcome of trial t-1 (Rew+, Rew-, or Miss). An additional switching analysis was conducted based on only the reward outcome of trial t-1 (i.e., rewarded versus non-rewarded trials) by collapsing Rew-and Miss trials together. One-sample *t*-tests were used to evaluate if differences in choice and switching biases deviated significantly from each other.

To further evaluate potential predictors of switching, a logistic regression was conducted using choice switching on trial t as the outcome variable (1 for switch, 0 for stay). Seven predictors were entered into the regression: 1) The reward outcome of trial t-1 (1 for reward, 0 for no reward), 2) the movement execution outcome of trial t-1 (1 for a hit, 0 for a miss), 3) the Rew-to Miss trial probability ratio of the chosen target on trial t, 4) the absolute cursor error magnitude on trial t-1 (distance from feedback cursor to target), 5) the veridicality of the feedback on trial t-1 (1 for veridical feedback, 0 for perturbed feedback), 6) the interaction of absolute error magnitude X the veridicality of the feedback on trial t-1, and 7) the current trial number. The multiple logistic regression was computed using the MATLAB function *glmfit*, with a logit link function. All regressors were normalized for display purposes. One-sample *t*-tests were used to test for significant regression weights across the sample. For two participants, full “separation” was observed with the reward regressor (e.g., they never switched after a Rew+ trial, or always switched after failing to receive a reward); these participants were excluded from the regression analysis, although they were included in all other analyses.

We also analyzed how movement feedback altered reaching behavior, in order to test whether participants were actively attempting to correct execution errors. In particular, we were interested in whether participants were sensitive to the non-veridical feedback provided on trials in which the feedback position of the cursor was perturbed. To assess this, we focused on trial pairs in which consecutive reaches were to the same target and the first trial of the pair was accurate (< ± 3° from target’s center), but the cursor feedback was displayed fully outside of the target, indicating a Miss (the analysis was conducted this way to limit simple effects of regression to the mean reaching angle). A linear regression was performed with the observed signed cursor error on the first trial of the pair as the predictor variable and the signed change in reach direction on the second trial as the outcome variable. One-sample *t*-tests were used to test for significant regression weights.

### Modeling analysis of choice behavior

A reinforcement-learning analysis was conducted to model participants’ choice data on a trial-by-trial basis and generate reward prediction error (RPE) time-courses for later fMRI analyses. We tested a series of temporal difference (TD) reinforcement-learning models (Sutton and Barto, 1998), all of which shared the same basic form:

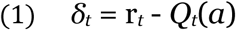

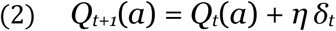

where the value (*Q*) of a given choice (*a*) on trial *t* is updated according to the reward prediction error (RPE) *δ* on that trial (the difference between the expected value *Q* and received reward r), with a learning rate or step-size parameter *η*. All models also included a decay parameter *γ* (Collins et al., 2014), which governed the decay of the three *Q*-values toward their initial value (assumed to be 1/the number of actions, or 1/3) on every trial:

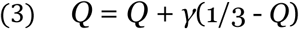

The decay parameter was important for model fitting, likely due to both the lack of any optimal slot machine and the stationary reward probabilities – many participants switched their choices often. Models without the decay parameter performed significantly worse than those with this parameter (data not shown).

Our previous results showed that participants discount Miss trials, suggesting a tendency to stay with a given choice following perceived execution errors (McDougle et al., 2016; Parvin et al., 2018) more often than they do following a choice error (Rew-trials). However, it is not known if this tendency is driven purely by RPE computations, or arises from a different source. To model two possible routes to “Miss discounting,” we included a persistence parameter, *Φ*, in the softmax computation of the probability of each choice (*P*),

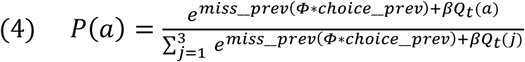

where “miss_prev” and “choice_prev” are indicator vectors, indicating, respectively, whether the previous trial was a Miss (1 for Miss, 0 for Rew+/Rew-) and which action was chosen, and *β* is the inverse temperature parameter. If *Φ* is positive, the learner is more likely to repeat the same choice after a Miss trial as a “bonus” of *Φ* is given to that option; if *Φ* is negative, the learner is more likely to switch after a Miss due to a “penalty” of *Φ*. This parameter represents a bias factor distinct from RPE-driven value updating (Bornstein et al., 2017) as the bonus (or penalty) is fixed regardless of the value of the chosen option.

We modeled reinforcement learning based on trial outcomes as follows: In the Standard(2η) model, distinct learning rates, *η*, were included to account for updating following negative RPEs (unrewarded trials) and positive RPEs (rewarded trials),

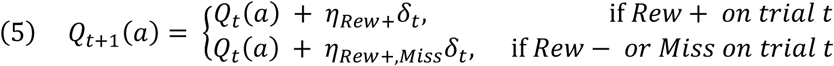

where *η_Rew+_* and *η_Miss/Rew-_* are the learning rates for updates following Rew+ or Miss/Rew-trials, respectively. Allowing positive and negative RPEs to update *Q* values at different rates has been shown to provide better fits to human behavior compared to models in which a single learning rate is applied after all trials (Gershman, 2015; Niv et al., 2012). We also included a second variant of this model, the Standard(no-*Φ*) model, that was identical to the Standard(2*η*) model but did not include the *Φ* parameter.

Two other models were included, based on our previous study in which negative outcomes could result from execution or selection errors (McDougle et al., 2016). One model, the Gating model, was similar to the Standard(2*η*) model, except that it had unique learning rates for each of the three possible trial outcomes (*η_Rew+_*, *η_Rew-_*, and *η_Miss_*). Thus, the Gating model allows for values to be updated at a different rate following execution errors (Miss) or selection errors (Rew-). Last, the Probability model separately tracked the probability of successful execution (*E*) for each target and the likelihood (*V*) of receiving a reward if execution was successful:

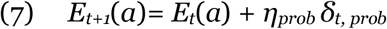

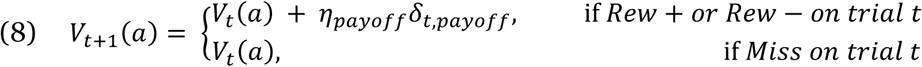

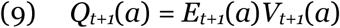

where *δ_t,prob_* and *δ_t,payoff_* represent, respectively, prediction errors for whether the current action was successfully executed (where r = 1 on Rew+/Rew-trials and r = 0 on Miss trials), and if a reward was received given that execution was successful.

Using the MATLAB function *fmincon*, all models were fit to each participant’s observed choices outcomes by finding the parameters that maximize the log posterior probability of the choice data given the model. To simulate action selection, *Q*-values in all models were converted to choice probabilities using a softmax logistic function (equation 4). All learning rate parameters (*η*) were constrained to be between −1 and 1. Negative values were permitted given that we did not have an *a priori* reason to assume *η_Miss_* would be positive, and thus opted to be consistent across all learning-rate parameters and models. The persistence parameter (*Φ*) was constrained to be between −5 and 5, and the decay parameter (*γ*) was constrained to be between 0 and 1. The temperature parameter (*β*) was constrained to be between 0 and 100, and a Gamma(2,3) prior distribution was used to discourage extreme values (Leong et al., 2017). *Q*-values for each target were initialized to 1/3.

The fitting procedure was conducted 100 times for each model using different randomized starting parameter values to avoid local minima during optimization, and the resulting best fit was used in further analyses. Model fits were evaluated using both the Bayesian information criterion (BIC; Schwarz, 1978) and Akaike information criteria (AIC; Akaike, 1974).

After model fitting and model comparison, we performed simulate-and-recover experiments on each of the four models to assess model confusability (Wilson et al., 2013). Choices were simulated for each model using the best-fit parameters of each of the 20 participants, yielding 20 simulations per model. Simulated data were then fit with each model (using 20 randomized vectors of starting parameters for each fit to avoid local minima) to test whether the correct models were recovered. Confusion matrices were created comparing differences in both individual and summed Aikake weights (Wagenmakers and Farrell, 2004), as well as the percent of simulations fit best by each model.

### fMRI data acquisition

Whole-brain imaging was conducted on a 3T Siemens PRISMA scanner, using a 64-channel head coil. MRI-optimized pillows were placed about the participant’s head to minimize head motion. At the start of the scanning session, structural images were collected using a high-resolution T1-weighted MPRAGE pulse sequence (1 × 1 × 1 mm voxel size). During task performance, functional images were collected using a gradient echo T2*-weighted EPI sequence with BOLD contrast (TR = 2000 ms, TE = 28 ms, flip angle = 90°, 3 × 3 × 3 mm voxel size; 36 interleaved axial slices). Moreover, a field map was acquired to improve registration and limit image distortion from field inhomogeneities (for one participant a field map was not collected).

Functional data were collected in a single run that lasted approximately 40 min. For one participant, the run was split into two parts due to a brief failure of the drawing tablet. Because of the self-paced nature of the reaching task (i.e., variable time taken to return to the start position for each trial, reach, etc.), the actual time of the run, and thus number of total TRs, varied across participants. The run was terminated once the participant had completed all 300 trials of the task.

### fMRI data analysis

Preprocessing and data analysis were performed using FSL v. 5.98 (FMRIB) and SPM12. Given the movement demands of the task and length of the scanning run, multiple steps were taken to assess and minimize movement artifacts. After manual skull-stripping using FSL’s brain extraction tool (*BET*), we performed standard preprocessing, registering the functional images to MNI coordinate space using a rigid-body affine transformation (*FLIRT*) applying the field map correction, spatially smoothing the functional data with a Gaussian kernel (8 mm FWHM), and attaining six column-wise realignment parameters derived from standard motion correction (*MCFLIRT*). To identify and remove components identified as head-motion artifacts, we then applied the independent components motion-correction algorithm ICA-AROMA (Pruim et al., 2015) to the functional data. As a final preprocessing step, we temporally filtered the data with a 100 s high-pass filter. Based on visual inspection of the data, four participants were excluded from further analyses, before preprocessing, due to excessive (> 3 mm pitch, roll, or yaw) head motion.

Four GLMs were performed. For the first three GLMs, we imposed a family-wise error cluster-corrected threshold of p < 0.05 (FSL FLAME 1), with a cluster-forming threshold of p < 0.001. Task-based regressors were convolved with the canonical hemodynamic response function (double Gamma), and the six motion parameters were included as regressors of no interest.

The first GLM was designed to functionally define ROIs that were sensitive to reward. Trial outcome regressors for the three trial types (Rew+, Rew-, Miss) were modeled as delta functions concurrent with visual presentation of the trial outcome. Task regressors of no interest included boxcar functions that spanned both the wait period and reach period. The contrast Rew+ > (Rew- and Miss) was performed to functionally identify reward-sensitive ROIs. Resulting ROIs were visualized, extracted, and binarized using the *xjview* package for SPM (http://www.alivelearn.net/xjview). Beta weights were extracted from the resulting ROIs using FSL’s *featquery* function. To identify areas sensitive to visuomotor errors while controlling for reward, we also tested a second trial outcome contrast: Miss > Rew-.

A second GLM was used to measure reward prediction errors (RPEs). Three separate parametric RPE regressors, corresponding to RPEs for each outcome, were entered into the GLM to account for variance in trial-by-trial activity not captured by the three binary outcome regressors (which were also included in the model). Beta weights for each RPE regressor were extracted from the striatum ROI (i.e., the functional “reward” ROI obtained from the first GLM) using FSL’s *featquery* function. Nuisance regressors included the wait period, reach period, and the three outcome regressors.

The third GLM was designed to identify brain areas parametrically sensitive to motor execution error magnitude. The regressor of interest here was limited to Miss trials and included a single separate parametric absolute cursor error regressor, which tracked the magnitude of angular cursor errors on Miss trials. Nuisance regressors included the wait period, reach period, and the three outcome regressors.

The fourth GLM was an exploratory psychophysical interaction (PPI) analysis (Friston et al., 1997). In a PPI, a task-specific regressor and ROI time course regressor are included in the same model with the critical addition of a third regressor that models the interaction between the other two regressors, capturing variance in activity not singularly attributable to either regressor alone. A mean time series from the striatum ROI was extracted using *fslmaths*, and added (unconvolved) to the model as an additional regressor. Interaction regressors between the striatum time course and the three individual outcome regressors were also included. Nuisance regressors included the wait period, reach period, and the three outcome regressors. We imposed a family-wise error cluster-corrected threshold of p < 0.05 (FSL FLAME 1), with a relaxed cluster-forming threshold of p < 0.05 (see Results).

All voxel locations are reported in MNI coordinates, and all results are displayed on the average MNI brain.

## Results

We developed a simple 3-arm “bandit task” in which, during fMRI scanning, the participant had to make a short reaching movement on a digital tablet to indicate their choice on each trial and to attempt to maximize monetary earnings (Figure 1). At the end of the movement, feedback was provided to indicate one of three outcomes, as follows: On Rew+ trials, the visual cursor landed in the selected stimulus and a money bag indicated that $.10 had been earned. On Rew-trials, the visual cursor landed in the selected stimulus but an X was superimposed over the money bag, indicating that no reward was earned. On Miss trials, the visual cursor was displayed outside the chosen stimulus (and no money was earned). The reward probability for each stimulus (“bandit”) was fixed at 0.4, but the probabilities of Rew- and Miss varied between the three stimuli (0.5/0.1, 0.3/0.3, 0.1/0.5 respectively; see Methods). Thus, we used a stationary multi-armed bandit task, as all probabilities were fixed.

### Choice Behavior

In previous studies using a similar task, participants showed a bias for stimuli in which unrewarded outcomes were associated with misses (execution errors) rather than expected payoffs (selection errors), even when the expected value for the choices were held equal (McDougle et al., 2016; Parvin et al., 2018). We hypothesized that this bias reflected a process whereby execution failures lead to attenuated negative prediction errors, with the assumption that “credit” for the negative outcome under such situations was attributed to factors unrelated to the intrinsic value of the chosen action.

In the current task, a similar bias could lead participants to prefer the high-Miss stimulus (0.5/0.1 ratio of Miss/Rew-outcome probabilities). However, the overall choice data showed only a weak bias across the three stimuli (Figure 2A, all *ps* > 0.15). We note that, unlike in our previous studies (McDougle et al., 2016; Parvin et al., 2018), the probability and magnitude of reward on each trial was identical for each stimulus.

**Figure 2:**
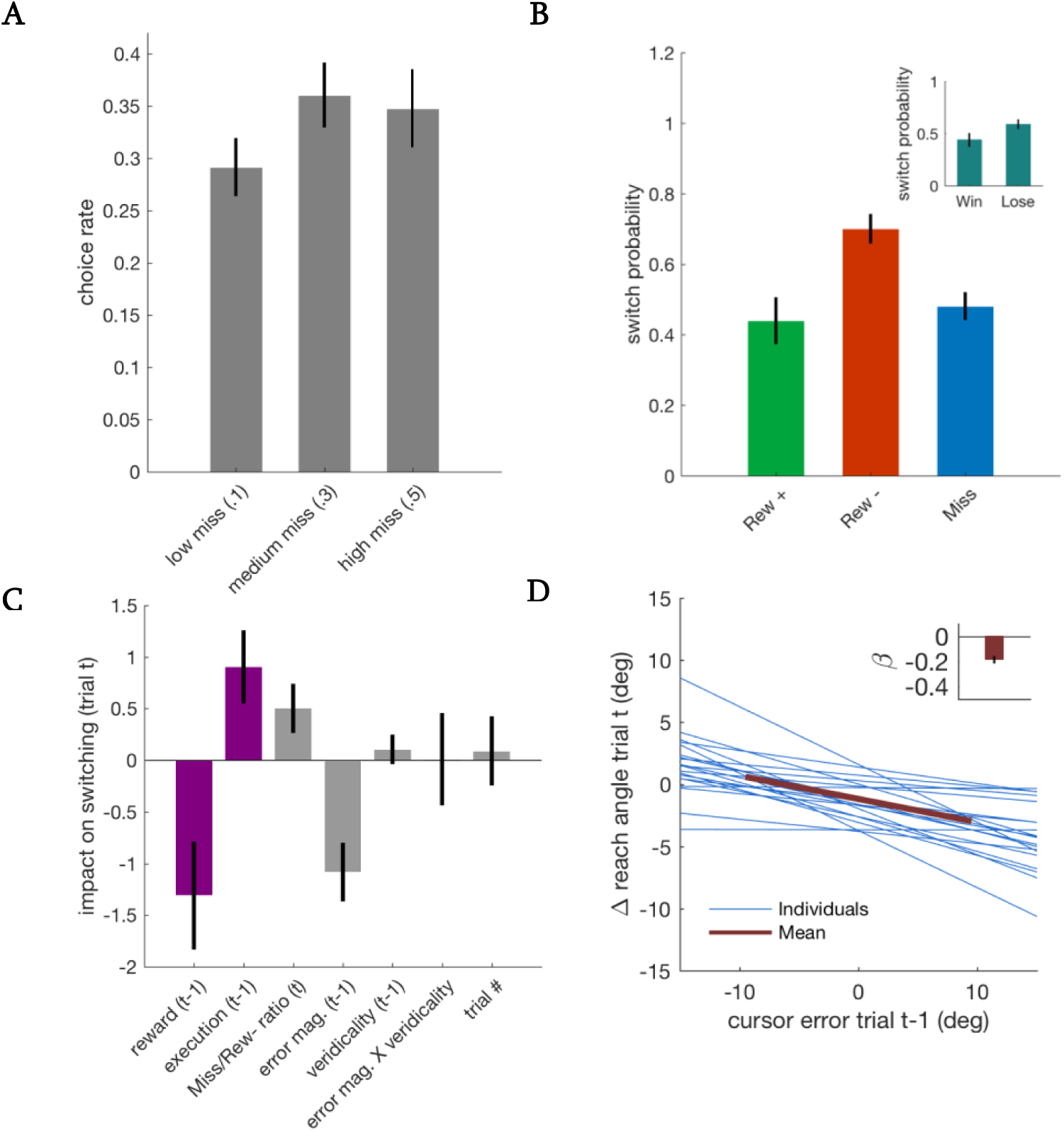
Behavior. **(A)** Participants’ biases to select stimuli with a different ratio of Rew-to Miss trials. **(B)** Average switch probabilities separated by the outcome on the previous trial. Inset: switch probabilities separated by rewarded trials (Rew+) versus unrewarded trials (Rew- and Miss, collapsed). **(C)** Logistic regression on switch behavior. **(D)** Logistic regression on change in reach angle as a function of signed cursor errors on the previous trial. This analysis is limited to trials in which participants' reach on trial t-1 was accurate, but the cursor was perturbed away from the target (Miss trial). Inset: average regression weight. Error bars = 1 s.e.m.

Critically, trial-by-trial switching behavior offers a more detailed way to look at choice biases (Figure 2B). Consistent with previous results, participants were more likely to switch to a different stimulus following Rew-trials compared to Miss trials (*t*_19_ = 5.08, *p* < 0.001). Moreover, they were more likely to switch after Rew-trials compared to Rew+ trials (*t*_19_ = 4.14, *p* < 0.001), and showed no difference in switching rate after Rew+ and Miss trials (*t*_19_ = 0.78, *p* = 0.45). Overall, participants were, on average, more likely to switch following a non-rewarded trial (Rew-or Miss) than a rewarded one (Rew+; *t*_19_ = 11.99, *p* < 0.001; Figure 2B inset), suggesting that they were generally sensitive to receiving a monetary reward, even though each lottery was identical for each slot machine. In sum, the switching behavior indicates that participants responded more negatively to Rew- outcomes compared to Miss outcomes, even though both yielded identical economic results. This finding is consistent with the hypothesis that cues suggesting a failure to properly implement a decision affect how value updates are computed.

A regression analysis was used to further probe switching behavior (Figure 2C). The first two regressors, reward and execution outcome, recapitulated the results shown in Figure 2B, where the reward outcome (reward vs. no reward) and the execution outcome (hitting the target vs. missing) both had a strong effect on switching behavior: Getting rewarded on trial t-1 negatively predicted switching on trial t (i.e., predicted staying over switching), reflecting the positive Rew+ trials (t-test for regression weight difference from 0: *t*_17_ = −2.38, *p* = 0.029;). In contrast, hitting the target on trial t-1 had a positive impact on the probability of switching on trial t, driven by the aversive Rew-trials (*t*_17_ = 2.42, *p* = 0.027;). Both effects were tempered by the Miss trials, which led to reduced switching (Figure 2B). Consistent with Figure 2A, the Rew-/Miss probability ratio of the selected target on trial t had only a marginal effect in the regression analysis (*t*_17_ = 2.01, *p* = 0.061).

Interestingly, the absolute magnitude of the cursor error on trial t-1 negatively predicted switching on trial t; that is, after relatively large errors, participants were more likely to repeat the same choice again (*t*_17_ = −3.62, *p* = 0.002). This effect did not appear to be driven by the veridicality of the error, as neither the regressor for the veridicality of feedback, nor the interaction between veridicality and error magnitude, predicted switching (*t*_17_ = 0.70, *p* = 0.49 and *t*_17_ = 0.02, *p* = 0.98, respectively). Lastly, switching behavior did not fluctuate over the duration of the experiment (“trial #” regressor; *t*_17_ = 0.26, *p* = 0.80).

### Effect of Feedback Perturbations

Perturbed cursor feedback was often required to achieve the desired outcome probabilities for each stimulus. Overall, we had to perturb the cursor position on 58.4% of trials. Most of these (47.6% of trials) were “false hits,” where the feedback cursor was moved into the target region following an actual miss. 10.8% of trials were false misses, in which the cursor was displayed outside the target following an actual hit.

We had designed the Miss-trial perturbations to balance the goal of keeping the participants unaware of the feedback perturbations, while also providing large, visually salient execution errors. The mean size of the perturbed Miss trial errors was 11.2° larger than veridical Miss trial errors (*t*_19_ = 35.19, *p* < 0.001), raising the possibility that participants could be made aware of the perturbations. The results from a post-experiment questionnaire were equivocal: When asked if the feedback was occasionally altered, the mean response on a 7-point scale was 4.3, where 1 is “Very confident cursor location was fully controlled by me,” and 7 is “Very confident cursor location was partially controlled by me.” However, it is not clear if the question itself biased participant’s answers, so further analyses were conducted.

As noted above, in terms of switching, the logistic regression analysis indicated that participants responded similarly to trials following veridical or perturbed cursor feedback (Figure 2C, negligible weights for variables related to veridicality of the feedback). We next examined if adjustments in reaching direction were responsive to non-veridical errors, as they would be expected to after veridical errors. To this end, we analyzed trial pairs in which the same stimulus was chosen on two consecutive trials where the first reach had been accurate but resulted in a false miss (mean number of pairs per participant = 18.4). If participants “believe” the perturbed feedback, the second movement should be shifted in the opposite direction of the preceding perturbation. We note that while we could perform the same analysis following veridical misses or perturbed hits, a shift would be expected simply from regression to the mean, whereas in this case, the hand would generally be shifting away from the mean. Consistent with this prediction, a regression analysis showed that heading direction did indeed shift by a fairly large amount in the opposite direction of the perturbation on the subsequent trial (*t*_19_ = −6.36, *p* < 0.001; Figure 2D). This could be interpreted as resulting from implicit sensorimotor adaptation, explicit adjustments in aiming, or both (Taylor et al., 2014). Taken together, both the regression and movement analyses, and to a lesser extent the questionnaire, indicate that manipulation of the cursor feedback did not have a significant impact on participants’ choice behavior (see Discussion).

### Modeling Results

We fit the participants’ trial-by-trial choice behavior with the four reinforcement learning models described in the Methods section (Figure 3). All models predicted trial-by-trial choice behavior better than chance (*t*-tests vs chance value of 0.33: all *p*′s < 0.001; Figure 3C). To perform a formal model comparison that considered the number of free parameters in each model, we calculated both the Bayesian (BIC) and Akaike (AIC) information criteria values for fits of each model (both metrics yielded similar results). First, the Gating model provided the best fit compared to the other three models in terms of both BIC and AIC (all *p′*s < 0.001, Figure 3A, B). Second, the Gating model had a higher average per-trial likelihood of predicting choices over the next best model (*t*_19_ = 4.61, *p* < 0.001; Figure 3C). Third, the Gating model provided the best fit for 16 of the 20 of the participants (Figure 3D). Consistent with our previous results (McDougle et al., 2016), the modeling analysis indicates that in tasks that allow for execution failures, an update parameter (*η*) devoted to such trials improves the model fit.

**Figure 3:**
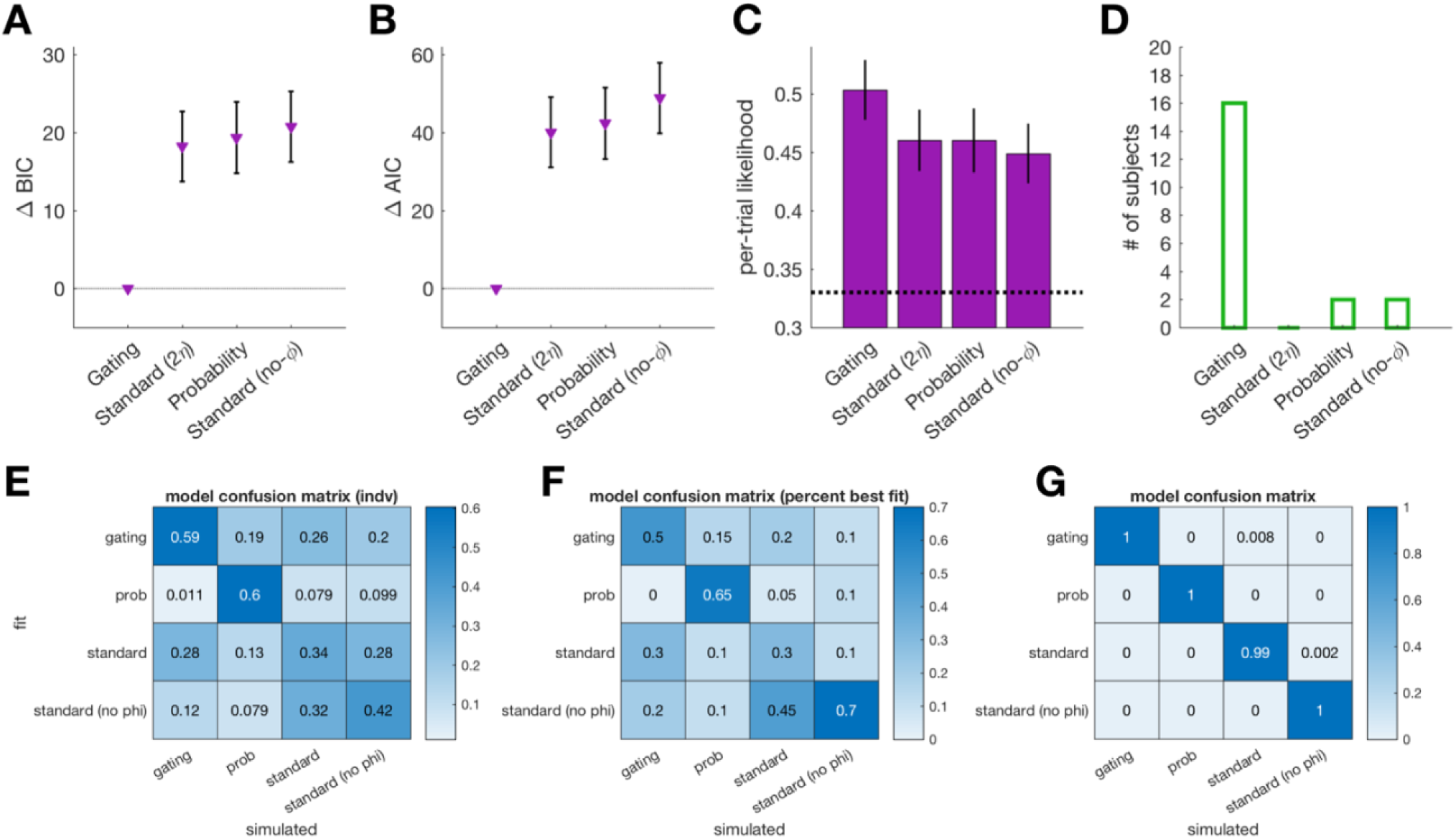
Model Comparisons. (**A**) Bayesian information criterion (BIC) and (**B)** Akaike information criterion (AIC) comparisons of each model. (**C**) Average per-trial likelihoods of each model predicting the participant’s true choice. (**D**) Number of participants best-fit by each model (using AIC). (**E-G**) Confusion matrices from the simulate-and-fit analysis, with the ground-truth simulated model on the x-axis and the model used to fit the simulation on the y-axis. Color indicates(**E**) average individual Akaike weights (an approximation of the conditional probability of one model over the others), (**F**) the percent of simulations best-fit by each model (using raw AIC values), and (**G**) summed Akaike weights across the sample. Error bars = 1 s.e.m.

We next examined the estimated parameter values for the Gating model. Parameter values were not normally distributed, and Wilcoxon sign-rank tests were thus used for statistical comparisons. The learning rates on Miss trials, *η_Miss_*, and Rew-trials, *η_Rew-_*, were both greater than zero (*p* = 0.010 and *p* = 0.014, respectively). The learning rate on Rew+ trials, *η_Rew+_* was marginally greater than zero (*p* = 0.09). As predicted, the *η_Miss_* parameter showed the lowest value (medians: *η_Miss_* = 0.07, *η_Rew+_* = 0.13, *η_Rew-_* = 0.23). However, a sign-rank test revealed no significant difference between *η_Miss_* and *η_Rew-_* (*p* = 0.18). Lastly, The persistence parameter (*Φ*) was significantly greater than zero (*p* = 0.023). This observation suggests that choice persistence after Miss trials may be driven by a top-down influence on action values during the choice phase.

Each model has several free parameters and they all share a similar form, raising a concern about model confusability. To address this, we simulated choice data with each model using its best-fit parameter values from each of the 20 participants, and then refit the simulations with each model (see Methods). If the models are reliably separable, each simulation should be best-fit by the model originally used to generate that simulation. The two models that best fit the behavioral data, Gating and Standard(2*η*), were modestly separable (Figure 3E, F), with respective average conditional probabilities of 0.59 versus 0.28 for fits to the Gating model simulations, and 0.26 versus 0.34 for fits to the Standard(2*η*) model simulations. We note that these values are the mean of each fit’s Akaike weight, which is an approximation of the model’s conditional probability versus the others (Wagenmakers and Farrell, 2004). As expected, the two Standard models were generally confusable with one another (Figure 3E, bottom right quadrant). The proportion of simulated agents from each model best fit by those same models is shown in Figure 3F. At the group level, summing AIC values over each full set of fits for each model (and computing Akaike weights on those sums) revealed rather strong model separability in all four cases (Figure 3G; we note, however, that summing tends to inflate differences in fit). Overall, this analysis suggests that the model fitting results should be interpreted with caution as each model is only subtly different. It is important to note that the primary reason modeling was conducted in the present study was to generate time courses of RPEs for the analysis of BOLD data. Indeed, the pattern of RPEs generated for each outcome (Rew+, Rew-, Miss) were very similar across models.

Previous studies have shown that movements toward high value choices are more vigorous (i.e., faster) compared to low value choices (Niv et al., 2007; Reppert et al., 2015; Seo et al., 2012). Given that we used reaching movements in the current study, we can ask if this phenomenon is observed in the current context, looking at the effect of model-derived *Q*-values on both reaction time (RT) and movement time (MT) on each trial. Overall, reaction times were moderately fast (µ = 0.59 ± .13 s) and movement times were quite fast (µ = 0.13 ± .06 s). These values, as well as the modeled *Q*-values of selected choices (from the Gating model), were extracted for each participant, de-trended using linear regression (due to gradual trends in both the RT and *Q*-value time courses), and then z-scored. Linear regressions were performed to quantify the influence of *Q*-values on trial-by-trial MTs and RTs. Consistent with previous results on movement vigor and value, *Q*-values negatively predicted MT (regression beta values relative to 0: *t*_19_ = −3.28, *p* = 0.004). In other words, higher-value choices were accompanied by faster movements (shorter movement times). No significant relationship was observed between RT and relative *Q*-values (*t*_19_= 0.38, *p* = 0.71). We speculate that this null result may be a function of the design of the task (Figure 2), which included an enforced wait period before movement. The MT result both agrees with previous research on vigor and value, and provides a case where our model describes behavioral data that were not part of the fitting procedure.

### Imaging

Figure 4A and Table 1 show the results of the whole-brain contrasts for reward processing (Rew+ > Rew- and Miss), and motor error processing (Miss > Rew). The reward contrast revealed four significant clusters spanning bilateral striatum, bilateral ventromedial prefrontal cortex (vmPFC), bilateral posterior cingulate (PCC), and a single cluster in left orbital frontal cortex (OFC). These ROIs are broadly consistent with areas commonly associated with reward (McClure et al., 2004; Schultz, 2015). For the motor error contrast, three broad clusters were revealed, including a single elongated cluster spanning bilateral premotor cortex (PMC), supplementary motor area (SMA), and the anterior division of the cingulate (ACC), as well as two distinct clusters in both the left and right inferior parietal lobule (IPL). This pattern is consistent with previous work on cortical responses to salient motor errors (Diedrichsen et al., 2005; Krakauer et al., 2004; Seidler et al., 2013).

**Table 1:** Significant Clusters. All clusters survived cluster correction at the p > 0.05 level (FLAME 1) with cluster-forming threshold of p < 0.001, with the exception of the PPI analysis, which used a threshold of p #x003C; 0.05. Coordinates are in MNI space and correspond to the cluster's center of gravity. vmPFC = ventromedial prefrontal cortex; OFC = orbitofrontal cortex; PCC = posterior cingulate cortex; SMA = supplementary motor area; ACC = anterior cingulate cortex; IPL = inferior parietal lobule; IFG = inferior frontal gyrus; M1 = primary motor cortex; PMC = premotor cortex; V1 = primary visual cortex; Cb = cerebellum; LOC = lateral occipital cortex.

| Analysis/Region | x (mm) | y (mm) | z (mm) | # voxels |
| --- | --- | --- | --- | --- |
| **Rew+ > (Rew- U Miss)** |  |  |  |  |
| striatum | 5 | 11 | -6 | 691 |
| vmPFC | 0 | 46 | 2 | 2710 |
| L OFC | -37 | 33 | -12 | 567 |
| PCC | -1 | -51 | 32 | 911 |
| **Miss > Rew-** |  |  |  |  |
| SMA/PMC/ACC | 10 | -2 | 61 | 1733 |
| R IPL | 60 | -24 | 33 | 1395 |
| L IPL | -55 | -25 | 26 | 733 |
| **PPI (Miss X Striatum)** |  |  |  |  |
| L IFG/L putamen | -40 | 20 | 2 | 944 |
| **Error Size (Miss)** |  |  |  |  |
| M1/PMC | 0 | -23 | 55 | 10399 |
| V1/R Cb | 15 | -80 | 9 | 2704 |
| R LOC/R IPL | -55 | -25 | 26 | 568 |

**Figure 4:**
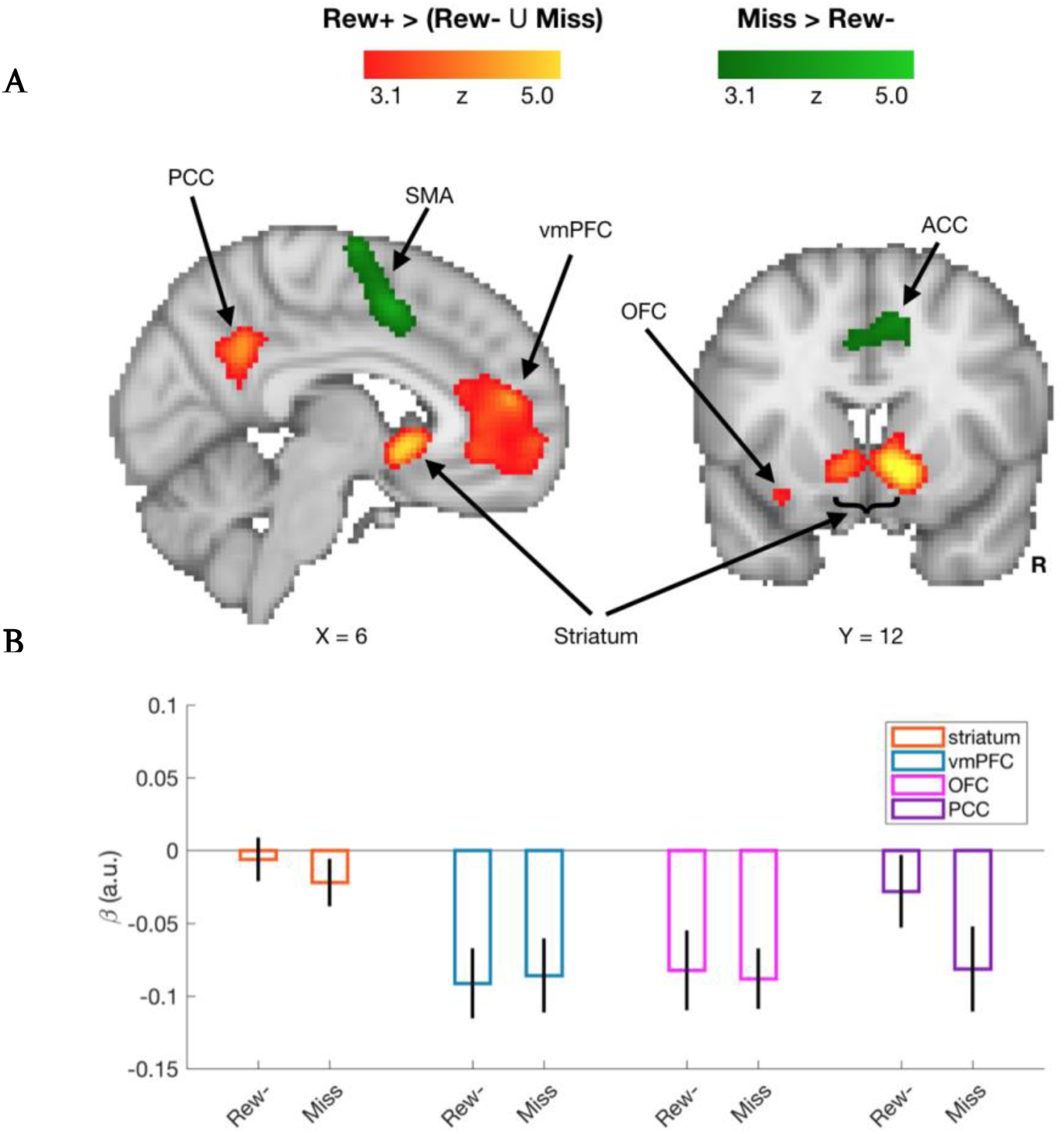
Trial Outcome Contrasts. (A) Results of whole-brain contrasts for Rew+ trials > Rew- and Miss trials (red/yellow), and Miss trials > Rew-trials (green). In the reward contrast (red/yellow), four significant clusters were revealed, in bilateral striatum, ventromedial prefrontal cortex (vmPFC), left orbital-frontal cortex (OFC), and posterior cingulate cortex (PCC). For the motor error contrast (green), three significant clusters were revealed, with a single cluster spanning bilateral premotor cortex, supplementary motor area (SMA), and the anterior division of the cingulate (ACC), as well as two distinct clusters in both the left and right inferior parietal lobule. (B): Beta weights extracted from each reward contrast ROI for the (orthogonal) Rew- and Miss trial outcomes. Error bars = 1 s.e.m.

Examination of feedback-locked betas on Rew- and Miss trials could identify gross differences in activity in these ROIs (Figure 4B), distinct from the more fine-grained parametric RPE modulations to be explored in the model-driven analysis (see below). Directly comparing the two negative outcome trial types revealed that average activity in the four ROIs was similar for Rew- and Miss trials, with no significant differences seen in the striatum (*t*_19_ = 0.88, *p* = 0.39), vmPFC (*t*_19_ = −0.24, *p* = 0.81), nor OFC (*t*_19_ =0.25, *p* = 0.81), and a marginal difference in the PCC (*t*_19_ = 1.95, *p* = 0.07).

In our second GLM, separate parametric RPE regressors for the three possible trial outcomes were constructed by convolving trial-by-trial RPE values derived from the Gating model with the canonical hemodynamic response function (HRF). Beta weights for the three regressors were then extracted from the striatum ROI delineated by the first GLM. As seen in Figure 5, striatal activity parametrically tracked trial-by-trial RPEs following Rew+ trials (*t*_19_ = 3.26, *p* = 0.004) and Rew-trials (*t*_19_ = 2.76, *p* = 0.013).

**Figure 5:**
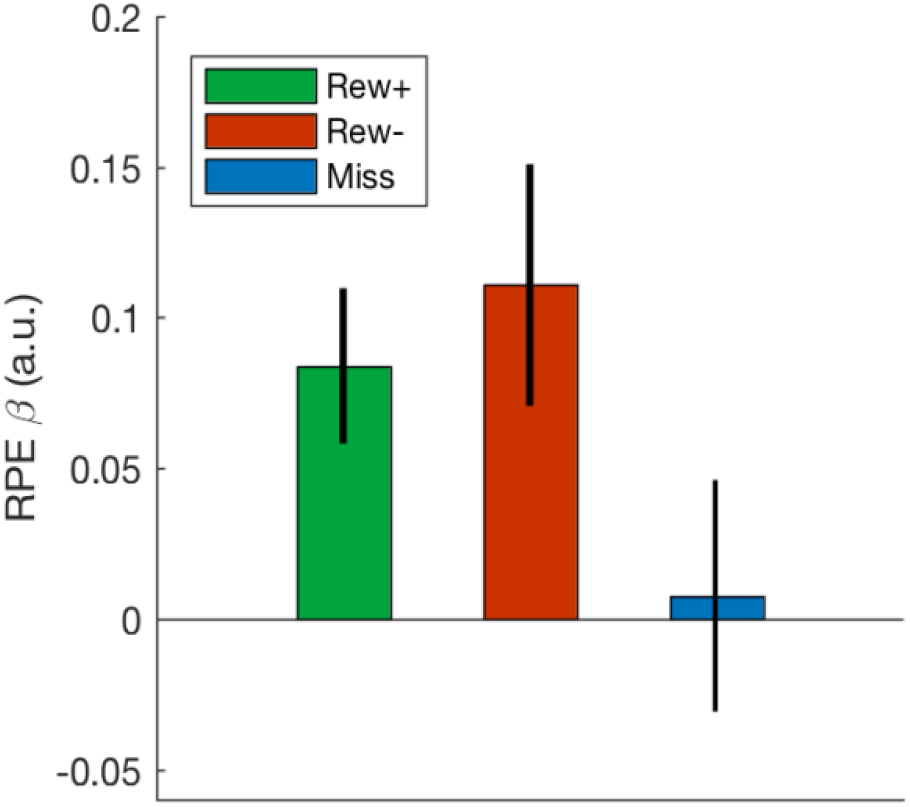
Outcome RPE coding in the striatum: Average reward prediction error (RPE) beta weights within the striatum ROI for each trial outcome type. Error bars = 1 s.e.m.

In contrast, striatal activity did not appear to encode RPEs following Miss trials (*t*_19_ = 0.20, *p* = 0.84). Critically, the strength of RPE coding was significantly greater on Rew-trials than on Miss trials (*t*_19_ = 2.52, *p* = 0.020), marginally greater on Rew+ trials than on Miss trials (*t*_19_ = 1.84, *p* = 0.082), and not significantly different between Rew+ and Rew-trials (*t*_19_ = −0.74, *p* = 0.47). Consistent with our hypothesis, these results suggest that striatal coding of RPEs is attenuated following execution failures. One consequence of this would be that choice value updating in the striatum would be effectively paused after miss trials, a strategy that could explain the observed behavioral biases (Figure 2B).

A third GLM analysis was conducted to confirm that the magnitude of observed execution errors was processed in predicted motor-related areas. This is distinct from the first GLM, which captured the effect of the mere presence of execution errors (Figure 4A, green). The absolute error size on Miss trials was entered as a parametric regressor in a whole brain analysis. Consistent with previous research (Anguera et al., 2009; Grafton et al., 2008), error magnitude was correlated with the modulation of activity in anterior cingulate cortex, dorsal premotor cortex, dorsal cerebellum (lobule VI), and primary visual cortex (Table 1). No significant voxels in the striatum were identified in this analysis, even at a relaxed cluster-forming threshold (*p* < 0.05).

To investigate areas that may act in concert with the ventral striatum in our task, we performed an exploratory psychophysiological interaction (PPI) connectivity analysis. Our PPI analysis quantifies correlations in BOLD activity between the striatal ROI and other brain areas that are more pronounced during Miss trials relative to the other two trial outcomes. Given the exploratory nature of the analysis and the conservative nature of PPIs, we relaxed our cluster-forming threshold to *p* < 0.05. The PPI revealed a significant functional interaction on Miss trials between the striatal ROI and an elongated cluster that consisted of, primarily, left inferior frontal gyrus (IFG) and left putamen (Figure 6).

**Figure 6:**
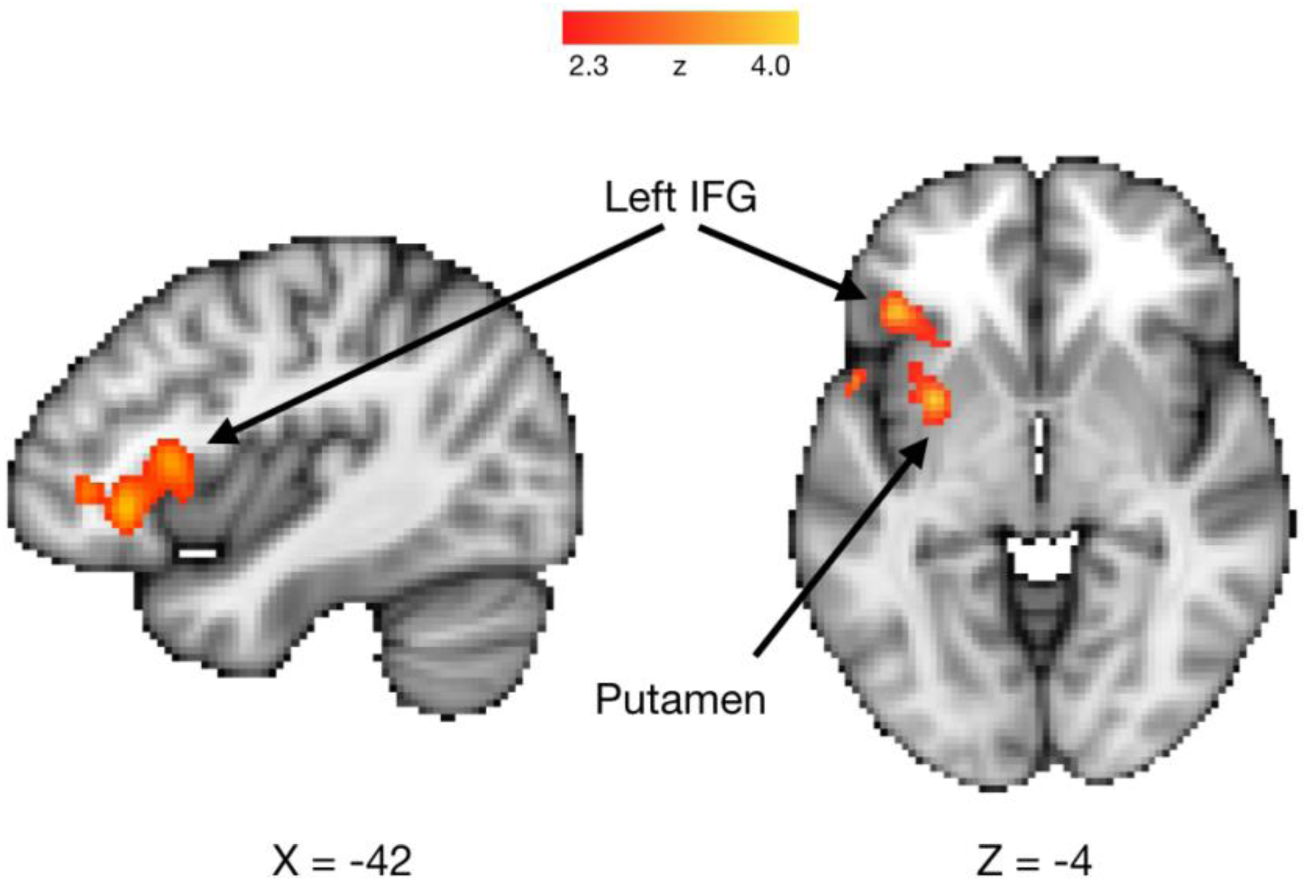
PPI Analysis. Activity in left inferior frontal gyrus (IFG) and left putamen was correlated with activity in the striatum ROI on Miss trials. Significant correlations were not found for Rew+ and Rew-trials.

As a point of comparison, we performed similar PPI analyses for both Rew+ and Rew-trials, comparing striatal connectivity in each versus the other two trial outcomes. Here, no significant clusters were found between the striatal ROI time course and the rest of the brain. One interpretation could by that because Rew+ and Rew-trials denote two sides of the same coin (standard reinforcement learning), these effects were washed out as connectivity patterns may be similar. We note that although the FSL FLAME algorithm used in our analyses limits false positive rate relative to most other approaches (Eklund et al., 2016), the clusters displayed in Figure 6 were not significant at more conservative statistical thresholds, and thus should be viewed with appropriate caution.

## Discussion

The present results demonstrate that perceived movement execution errors influence reward prediction error (RPE) computations in the human striatum. When participants did not receive a reward but properly executed their decision, the striatum predictably represented the corresponding negative RPE, consistent with much previous experimental work. However, on trials where a no-reward outcome was framed as the result of an action execution failure, the striatum did not appear to generate a corresponding negative RPE (Figure 5). These results indicate that before critiquing the quality of a decision, the striatum may use knowledge concerning whether the decision was properly implemented in the first place. This contingency was reliably observed in participants’ choice behavior (Figure 2), and can be described by a reinforcement learning model where decision execution errors demand a unique learning rate parameter (Figure 3).

These findings fit into a broader reevaluation of the nature of RPEs in the mesostriatal dopamine system. Mounting evidence suggests that the striatum does not just signal a model-free prediction error, but is affected by high-level cognitive states, concerning, for instance, model-based predictions of future rewards (Daw et al., 2011), sampling from episodic memory (Bornstein et al., 2017), top-down attention to relevant task dimensions (Leong et al., 2017), and the holding of stimulus-response relationships in working memory (Collins et al., 2017). We believe the present results add to this body of evidence, showing that contextual cues concerning the implementation of a decision affect if and how the represented value of that decision is updated by a prediction error.

We note that the putative “gating” phenomenon, the diminished encoding of a negative RPE in the striatum, was not categorical; indeed, participants displayed varying degrees of gating both behaviorally and neurally (Figure 2, Figure 5). One speculation could be that gating is a function of how optimistic a participant is that they could correct a motor error in the future. By this hypothesis, gating is useful only if one is confident in their execution ability, and are thus likely to persist with a decision until successful execution will allow them to glean reward information about the selected stimulus. On the other hand, if one is not confident in their ability to execute a movement, a negative RPE might also be generated upon an execution error, steering them away from that choice and its associated action in the future.

This hypothesis could explain a curious result in a previous study (McDougle et al., 2016): We found that participants with degeneration of the cerebellum, which results in problems with both motor learning and motor execution, showed diminished “gating” behavior; that is, they avoided decisions that were difficult to execute, even at the cost of larger rewards. We had hypothesized that the cerebellum may be an important structure in a putative gating mechanism, perhaps communicating sensory prediction errors to the basal ganglia via established bidirectional connections (Bostan et al., 2013). However, significant cerebellar activity only survived statistical correction in our analysis of cursor error size (Table 1), and the results of our planned analyses on trial outcomes did not reveal significant interactions between the cerebellum and striatum arguing against a cerebellar-dependent gating process. Indeed, a recent behavioral follow-up to our previous results suggests that cerebellar error signals are likely not affecting choice behavior in this kind of task (Parvin et al., 2018); rather, participants’ likely use some form of internal model concerning the causal structure of the task to guide their decisions (Green et al., 2010). It would be reasonable to assume that individuals with cerebellar degeneration may have a greater propensity to avoid choices associated with high execution errors given their reduced confidence in their ability to successfully control their movements.

Via reverse inference, the results of our connectivity analysis (Figure 6) suggest that the left inferior frontal gyrus (IFG) is one candidate region involved in the attenuation of RPEs following movement execution errors. Recent work suggests that the left IFG inhibits belief updating following certain negative outcomes (Moutsiana et al., 2015; Sharot et al., 2011, 2012), findings that are intriguingly similar to the results presented here. Others have highlighted a more general role for the left IFG in controlled retrieval processes that apply goal-relevant knowledge in a top-down fashion (Badre and Wagner, 2007). We speculate that a perceived execution error could be interpreted as a specific case of a more generalized cue about the current “state” the participant is in, where the specific implication of this putatively negative outcome is to inhibit value updating.

Although we are interpreting the current results in the context of perceived motor execution errors, an alternative explanation is that participants did not fully believe the feedback they received because it was often perturbed (see Methods). Thus, participants may have estimated whether they truly “caused” an observed outcome, and the gating of striatal RPEs may reflect instances where participants feel the outcome was manipulated. The power of each trial type by feedback veridicality/non-veridicality was too low across the group to test this hypothesis using a GLM on the imaging data (e.g., as few as 14 trials). However, we note that the most common perturbed-feedback trials involved situations in which the feedback was adjusted to hit the target (where the actual movement had missed the target), and, overall, Rew+ and Rew-trials showed robust RPE coding in the striatum (Figure 5). Moreover, the behavioral results suggest that error veridicality was not a strong predictor of participants’ choices (Figure 2C), nor movement kinematics (Figure 2D). Either way, future research should test the specificity of our results. For example, would the observed attenuation of RPEs happen if the lack of reward was clearly attributed to an external cause, for instance if the participant’s hand was knocked away by an external force? The results observed in the present study could reflect a unique role of intrinsically-sourced motor execution errors in RPE computations, or a more general effect of any arbitrary execution failure, whether internally or externally generated.

Research concerning the computational details of instrumental learning has progressed rapidly in recent years, and the nature of one fundamental computation in learning, reward prediction error, has been shown to be more complex than previously believed. Our results suggest that prediction errors update decisions in a manner that incorporates the successful implementation of those decisions, specifically, by ceasing to update value representations when a salient execution failure occurs. These results may add to our understanding of how reinforcement learning proceeds in more naturalistic settings, where successful action execution is often not trivial.

